# Understanding the adaptive evolution of mitochondrial genomes in intertidal chitons

**DOI:** 10.1101/2020.03.06.980664

**Authors:** Dipanjana Dhar, Debayan Dey, Soumalee Basu, Helena Fortunato

## Abstract

Mitochondria are the centre of energy metabolism in eukaryotic cells and its genes are thus key to the evolution of molecular mechanisms that metabolize cellular energy. Intertidal zone is one of the most stressful environments with extreme shifts in temperature, salinity, pH and oxygen concentrations. Marine molluscs, particularly chitons belong to the ecologically dominant organisms in this extreme environment, symbolizing an ideal model to understand mitochondrial stress adaptation. Here, we used concatenated mitochondrial genetic components separately from seven chitons of the intertidal zone to reconstruct phylogenetic relationships among these species. We performed selection analyses considering sites and branches of individual protein-coding genes to identify potentially adaptive residues and localize them in the protein structures of mt subunits. Our results exhibited significant amino acid changes in sites under diversifying selection of all the protein-coding genes, indicative of the adaptive evolution of mitochondrial genome in chitons. Furthermore, we obtained sites in the transmembrane helices lining the proton translocation channel as well as in surrounding loop regions, providing implication towards functional modification of the OXPHOS proteins essential for survival in dynamic environment of the intertidal zone.

## Introduction

Animal mitochondrial DNA is generally about 15,000-20,000 base pairs in size. However, much larger mitochondrial genomes are copies of duplicated genes rather than genetic variation in mtDNA **[1, 2]**. Mitochondrial DNA contains 13 protein-coding genes, 2 *r*RNAs of the mitochondrial ribosome and 22 *t*RNAs essential for translating OXPHOS proteins. Furthermore, presence of non-coding region(s) in the mtDNA is probably the site for controlling transcription and replication **[3]**. The 13 mitochondrial PCGs forming subunits of respiratory chain complexes: cytochrome *c* oxidase subunit 1-3 (*COX1, COX2, COX3*), cytochrome *b* (*CYTB*), NADH dehydrogenase subunit 1-6 (*ND1, ND2, ND3, ND4, ND5, ND6*), NADH dehydrogenase subunit 4L (*ND4L*), ATPase F0 subunit 6 (*ATP6*) and ATPase F0 subunit 8 (*ATP8*), are responsible for producing up to 95% of the energy of eukaryotic cells **[4]**. Complex I (*ND1, ND2, ND3, ND4, ND4L, ND5* and *ND6*), III (*CYTB*), IV (*COX1, COX2* and *COX3*) and V (*ATP6* and *ATP8*) are four of the five complexes required in the electron transport chain system and generation of ATP, combined with other subunits encoded by the nuclear DNA genome **[5]**.

Mitochondrial stress can have major consequences on the metabolic performance of an organism. Since mitochondrial proteins and mitochondria-derived stress signals regulate both oxidative phosphorylation and innate immune response, maintenance of the mitochondrial integrity and signaling therein are important for cellular homeostasis and survival **[6-11]**. All the events resulting to the impairment of mitochondrial functions have adverse effects on the amino acid constituency of the OXPHOS proteins. Changes in these amino acid residues do not alter protein structure rather can have an impact on the functional domains, such as regions lining the proton translocation channel and subunit interacting sites, and thereby succour animals to adapt to challenging environments **[12, 13]**. Therefore, estimation of selection pressures acting on mt proteins could extend deep insight into the adaptive evolution of mitochondrial genome. Most of the researches have been conducted on the adaptive evolution of mitochondrial genome in vertebrates, specially mammals and fish **[14, 15]**. However, fundamental questions regarding mitogenome evolution of invertebrates, particularly marine invertebrates, remain largely unanswered.

Mollusca is the second largest animal phyla after arthropods consisting of eight classes: Bivalvia, Cephalopoda, Gastropoda, Monoplacophora, Polyplacophora, Scaphopoda, Caudofoveata and Solenogastres, as per the widely accepted “Aculifera hypothesis” **[16-18]**. Among all the classes, chitons (Polyplacophorans) are the most primitive marine molluscs with fossil record tracing back to the early Cambrian **[19, 20]**. In addition, this phylum constitutes a ubiquitous, heterogeneous and economically important group of invertebrates that function as ecosystem engineers **[21]**. Although the most conspicuous symptom of any ecosystem deterioration is the decline or disappearance of sensitive species, molluscs display an unusual resilience towards environmental changes **[22]**. Chitons, mostly inhabitants of the intertidal zones, are competent enough to survive and maintain mitochondrial homeostasis regardless of regular oscillations of immersion and emersion, extreme alternations of temperature, salinity, pH and hydrodynamic forces in their surrounding environment **[23-25]**. More detailed studies on mitochondrial genome evolution will allow conducting further analysis on metabolic adaptation of intertidal chitons.

In spite of possessing significant roles as a global food resource and marine bioindicator, only a small percentage of molluscs possess completely sequenced and annotated genomes, rendering the phylum with the lowest ratio between fraction of sequenced genomes and number of described species **[26]**. At the time of writing this article (December 2019), complete mt genome sequences of only 7 chiton species were available on GenBank **[27-30]**. We therefore considered seven mt genomes representing species of the order Chitonida from the class Polyplacophora to determine evolutionary selection acting on the 13 mitochondrial protein-coding genes. We performed site and branch-site specific likelihood analyses to quantify the probability of positive selection on each site and branch of individual genes across all the seven chitons. Thereafter, we assessed changes in amino acids based on their physicochemical properties and mapped the positively selected sites in structure-based homology models to ascertain the effect of these changes in functionally important regions of the OXPHOS proteins. This work also focused on the reconstruction of phylogenetic relationships considering genetic components (whole genome, PCGs, *t*RNAs and *r*RNAs) of the mitochondrial DNA, separately. We also observed interspecific variations in the control region of the mitogenome of these polyplacophoran species, thus enhancing the existing knowledge on the mitogenomics of chitons.

## Materials and Methods

### Sequence retrieval

Complete mitochondrial genome sequences of seven Polyplacophoran species were downloaded from the National Center for Biotechnology Information (NCBI) database (https://www.ncbi.nlm.nih.gov/nucleotide/) **[31]**. The nucleotide and amino acid sequences of 13 mitochondrial PCGs for the same were also retrieved from the above mentioned database. Additionally, sequences pertaining to 2 *r*RNAs, 22 *t*RNAs and the non-coding control region were separately accumulated from the mitogenome of chitons. The details of the species considered for this study are provided in **Supp. Table 1**.

### Sequence alignment & phylogenetic analysis

Four nucleotide sequence datasets were constructed with the mitogenomes obtained from 7 chiton species – (A) 13 mitochondrial protein-coding genes (*COX1, COX2, COX3, CYTB, ATP6, ATP8, ND1, ND2, ND3, ND4, ND4L, ND5* and *ND6*); (B) 22 *t*RNAs; (C) 2 *r*RNAs and (D) highly variable control region. A species from the class Gastropoda (*Conus textile*, accession ID: DQ862058.1) was chosen as outgroup. For each of the aforementioned datasets, codon-based (for PCGs) and multiple sequence alignments were performed using MUSCLE **[32]** integrated in the MEGA-X software **[33]**. Besides, the entire mitogenomes were aligned in the Geneious Prime Version 2020.0.2 software (trial version), available at http://www.geneious.com. The aligned coding sequences for mitochondrial PCGs, *t*RNAs and *r*RNAs were then separately concatenated in MEGA-X, with default settings. The ambiguously aligned regions for individual alignments were removed using Gblocks server v0.91b **[34]**, prior to concatenation.

The appropriate nucleotide substitution models for each dataset were determined in jModelTest2 (v2.1.10), based on different information criterion calculations, starting with 11 substitution schemes and fixed BIONJ-JC option base tree for likelihood calculations **[35]**. We selected models according to Akaike Information Criterion (AIC) and corrected Akaike Information Criterion (cAIC). The best fit models obtained for whole mitogenome, PCGs, *t*RNAs, *r*RNAs and control region are TIM1+G, TVM+I+G, GTR+G, TVM+I+G and GTR respectively. Since most of the models obtained could not be implemented in MrBayes 3.2.6, they were substituted with General Time Reversible (GTR) model along with a proportion of invariable sites (I) and heterogeneity of substitution rates among sites modeled following a gamma distribution (G), wherever applicable **[36, 37]**. In addition, substitution saturation was calculated on each codon position of our dataset by Xia *et al*. statistical test **[38]**, executed in DAMBE software v7 **[39]**. Subsequently, the maximum log likelihood values for substitution matrix and transition/transversion rate ratio were calculated for the coding and non-coding regions of the mitogenome in seven chitons using MEGA-X.

Phylogenetic analyses, for each dataset, were performed using Bayesian inference with the aforementioned nucleotide substitution models. Phylogenetic trees were contrived using MrBayes v3.2.6 plug-in in the Geneious Prime Version 2020.0.2 software (trial version), via the CIPRES REST API **[40, 41]**. A Markov Chain Monte Carlo (MCMC) search with 5 million generations was performed, logging results every 500^th^ generation. The first 25% of the trees were discarded as burn-in. A consensus tree and the Bayesian Posterior Probabilities (BPP) were estimated based on the remaining trees. All the derived phylogenies generated were visualized using Figtree v1.4.3 (http://tree.bio.ed.ac.uk/software/figtree/) and iTOL v5 **[42]**.

### Selection Pressure Analyses

Estimation of ω *i*.*e*. the ratio of non-synonymous substitutions per non-synonymous sites (d_N_) to synonymous substitutions per synonymous sites (d_S_) indicate natural selection on protein coding genes. Under neutrality, the ratio is not expected to deviate significantly from 1 (ω = 1), whereas notable changes in the ratio may be interpreted in case of positive selection (ω>1) or negative selection (ω<1). The ω value was determined using the codon-based maximum likelihood (CODEML) algorithm implemented in EasyCodeML v1.21 **[43, 44]**. Both site and branch-site models were used to identify the variation of selective pressures on individual mitochondrial PCGs of the chitons. Seven codon substitution models described as M0 (one-ratio), M1a (nearly neutral), M2a (positive selection), M3 (discrete), M7 (beta), M8 (beta and ω>1) and M8a (beta and ω=1) were investigated and four likelihood ratio tests (M0 vs. M3, M1a vs. M2a, M7 vs. M8 and M8a vs. M8) were performed to measure the likelihood of positive selection on each site for individual PCGs **[45-50]**. On the other hand, branch-site models enable ω to differ among sites as well as across branches (foreground-lineages) of the phylogenetic tree **[51]**. Thus, we tested for positive selection on each branch of the phylogeny, considering a single foreground branch at a time. Bayes Empirical Bayes analysis was used to compute posterior probabilities in order to recognize sites under positive selection on the selected branches if the LRT *p*-values are significantly correct.

Additionally, we used the HyPhy package v2.5 (available from www.hyphy.org) to assess codons under selection pressure **[52]**. MEME (Mixed Effects Model of Evolution) and FEL (Fixed Effects Likelihood) analyses were carried out to detect individual sites subjected to episodic diversifying selection and purifying selection, respectively **[53, 54]**. Both the analyses were inferred at *p*-value<0.05 significance level. Furthermore, the impact of potential positive selection was detected using TREESAAP software v3.2, which detects changes in local physicochemical properties due to amino acid replacements **[55, 56]**. A sliding window equal to 20 codons alongwith magnitude categories ≥ 6 and *p*-values < 0.001 (z-score > 3.09) significance level were considered for accurate assessment of amino acid property changes.

### Protein modelling and structural analysis

In order to understand the influence of sites undergoing positive selection or exhibiting physicochemical property changes, we mapped them onto the three-dimensional structure of mitochondrial proteins. Owing to the unavailability of 3D structures of mt-subunits for molluscs in Protein Data Bank (PDB) **[57]**, we modeled the same for the Eastern beaded chiton *C*.*apiculata*. This species was selected among the seven chitons due to its ancestral mt genome architecture that presumably indicate the plesiomorphic condition for the phylum Mollusca. The structures of *COX* subunits and *CYTB* protein were modeled following significant similarities with templates sequences (PDB ID: 3ABM and 3H1I.1.C, respectively) using the SWISS-MODEL server **[58]**. The models for remaining mitochondrial proteins were generated in I-TASSER server, using default parameters **[59, 60]**. The overall structure of mitochondrial complex I and IV for *C*.*apiculata* were generated by structurally aligning the component subunits to corresponding complex I and IV templates from *Ovis aries* (PDB ID: 5LNK) and *Bos taurus* (PDB ID: 3ABM) respectively, using PyMOL software **[61]**. Additional information about the orientation of transmembrane domains of these proteins was obtained using the TMHMM server v2.0 **[62]**. PyMOL was further used for graphic modifications, visualization and depicting final illustrations.

## Results & Discussions

### Phylogenetic analyses & diversity

Based on the concatenated sequences of 13 mitochondrial PCGs, phylogenetic relationship among the polyplacophorans **(Fig. 1)** predominantly resembled the tree obtained by Guerra *et al*, 2018. The phylogenetic trees revealed family-wise clading of the chiton species with strong nodal support where families Mopallidae and Lepidochitonidae emerged sister to Chaetopleuridae and Chitonidae. However, the phylogeny derived from the whole mitogenomes of 7 species displayed contrasting topology **(Fig. 2)**, most probably owing to the alternative gene arrangement in *Sypharochiton sp*. This observation appeared highly congruent with the fact that family-specific rearrangements took place in Chitonidae due to inversion of *t*RNA_Val_ and *t*RNA_Trp_ genes. Besides, other members exhibited alterations in their *t*RNA arrangement as follows: (1) inversion of *t*RNA cluster (*t*RNA_Met_, *t*RNA_Cys_, *t*RNA_Tyr_, *t*RNA_Trp_, *t*RNA_Gln_, *t*RNA_Gly_ and *t*RNA_Glu_) in *C. caverna* and *N. californica*; (2) reversal of *t*RNA_His_ in *C. stelleri* and (3) transposition of *t*RNA_Asp_ particular to *K. tunicata*. It is worth mentioning that the clustering pattern of chitons in the phylogeny contrived using whole mitochondrial genome is reflective of the arrangement of their protein-coding genes and *t*RNAs **(Fig. 3)**.

**Figure 1:**
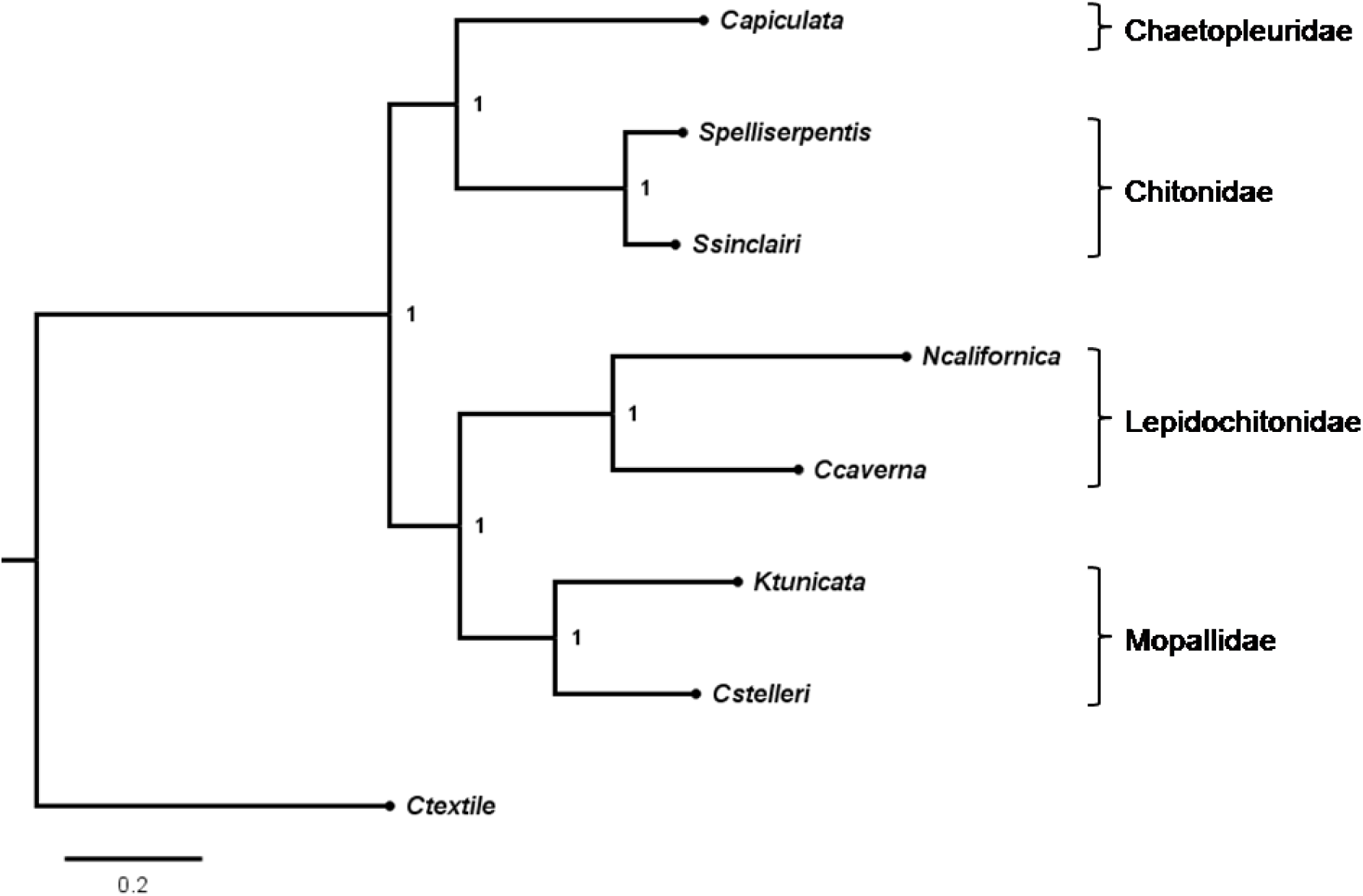
Consensus Bayesian phylogenetic tree based on the concatenated mitochondrial protein-coding genes from seven chitons. Posterior probabilities supporting each node are shown. *C. textile* (Gastropoda) was taken as outgroup. Families of the polyplacophorans corresponding to each clade are mentioned on the right side of the tree.

**Figure 2:**
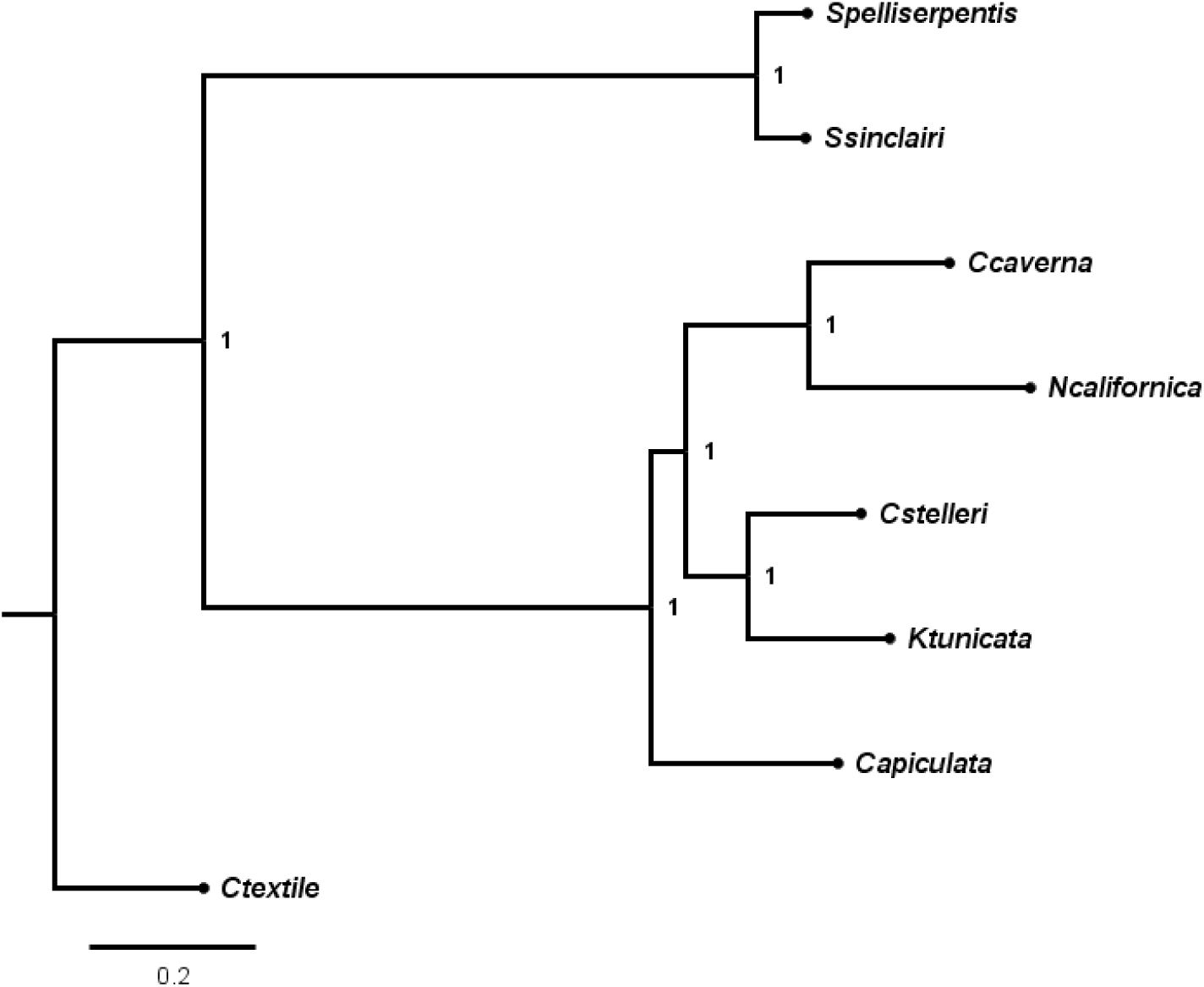
Bayesian phylogeny considering complete mitochondrial genome sequences of seven chitons. Posterior probabilities supporting each node are shown. *C. textile* (Gastropoda) was taken as outgroup.

**Figure 3:**
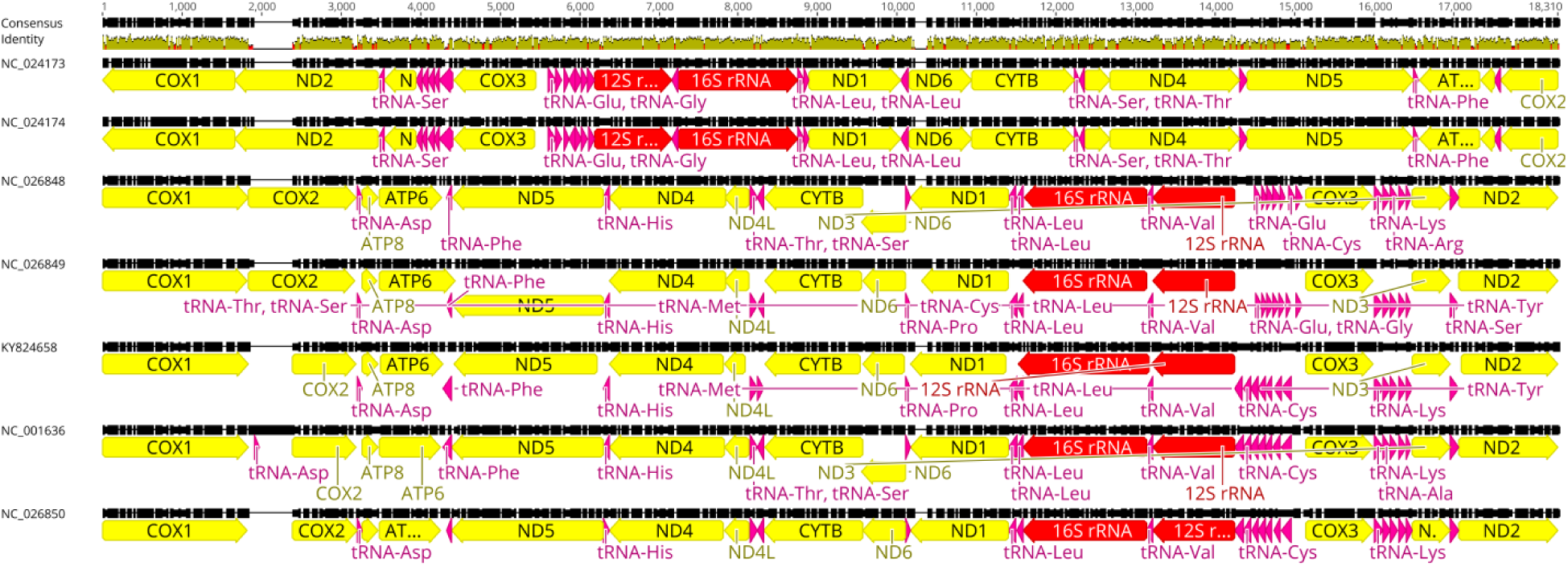
Schematic representation depicting arrangement of genetic components (PCGs, *t*RNAs & *r*RNAs) of the mitogenome in the seven Polyplacophoran species. The accession IDs in the figure are as follows-NC_024173.1 (*Sypharochiton sinclairi*), NC_024174.1 (*Sypharochiton pelliserpentis*), NC_026848.1 (*Cyanoplax caverna*), NC_026849.1 (*Nuttallina californica*), KY824658.1 (*Chaetopleura apiculata*), NC_001636.1 (*Katharina tunicata*) and NC_026850.1 (*Cryptochiton stelleri*).

Even though the entire mt-DNA remains linked and gets inherited as a single entity in the absence of recombination, coding and non-coding segments therein demonstrate differential substitution rates with higher selective forces acting on the functionally conserved protein-coding genes **[63]**. Therefore, we compared individual mitochondrial genetic components under varying selection pressure through phylogenetic clustering. Phylogeny inferred separately from concatenated nucleotide sequences of 22 *t*RNAs and 2 *r*RNAs **(Supp. Fig. 1a & 1b)** demonstrated identical clading pattern with the one contrived using PCGs. It can be stated that when all the *t*RNAs were taken into account, the phylogenetic relationship among the species was in accordance with other coding portions, regardless of their different arrangements in the mitogenome.

Similarly, we analyzed the highly polymorphic non-coding sequence (also called the control region) of the mitochondrial genome. This region is present between *t*RNA_Glu_ and *COX3* in all the chitons studied here, excepting two members in the Lepidochitonidae family where the region is shifted between *t*RNA_Glu_ and *12S r*RNA. Upon multiple sequence alignment **(Fig. 4)** we observed high sequence variability among these AT-rich regions, rendering it unsuitable for inferring spatial relationships from the control region alone. Thus, in spite of the utility of this mitogenomic part as a genetic marker for phylogenetic analysis in several metazoans **[64-66]**, the same is not applicable for polyplacophoran species considered here. Furthermore, we inferred incidences of nucleotide substitutions in the protein-coding genes and control region of chiton mitogenomes. Calculating transition/transversion bias (*R*) is important for deciphering genome evolution since it allows verifying the presence of bias in nucleotide conversions. In our datasets, transitions were more favoured over transversions **(Fig. 5)**. The maximum likelihood estimate of substitution matrix is depicted in **Supp. Table. 2**. Moreover, our results revealed the presence of more transitions leading to synonymous substitutions in protein-coding genes so as to maintain functional constancy in the three-dimensional structures **[67]**. In case of the non-coding control region, though transversions seemed to be prevalent, high transitions were retained to ensure proper regulatory function.

**Figure 4:**
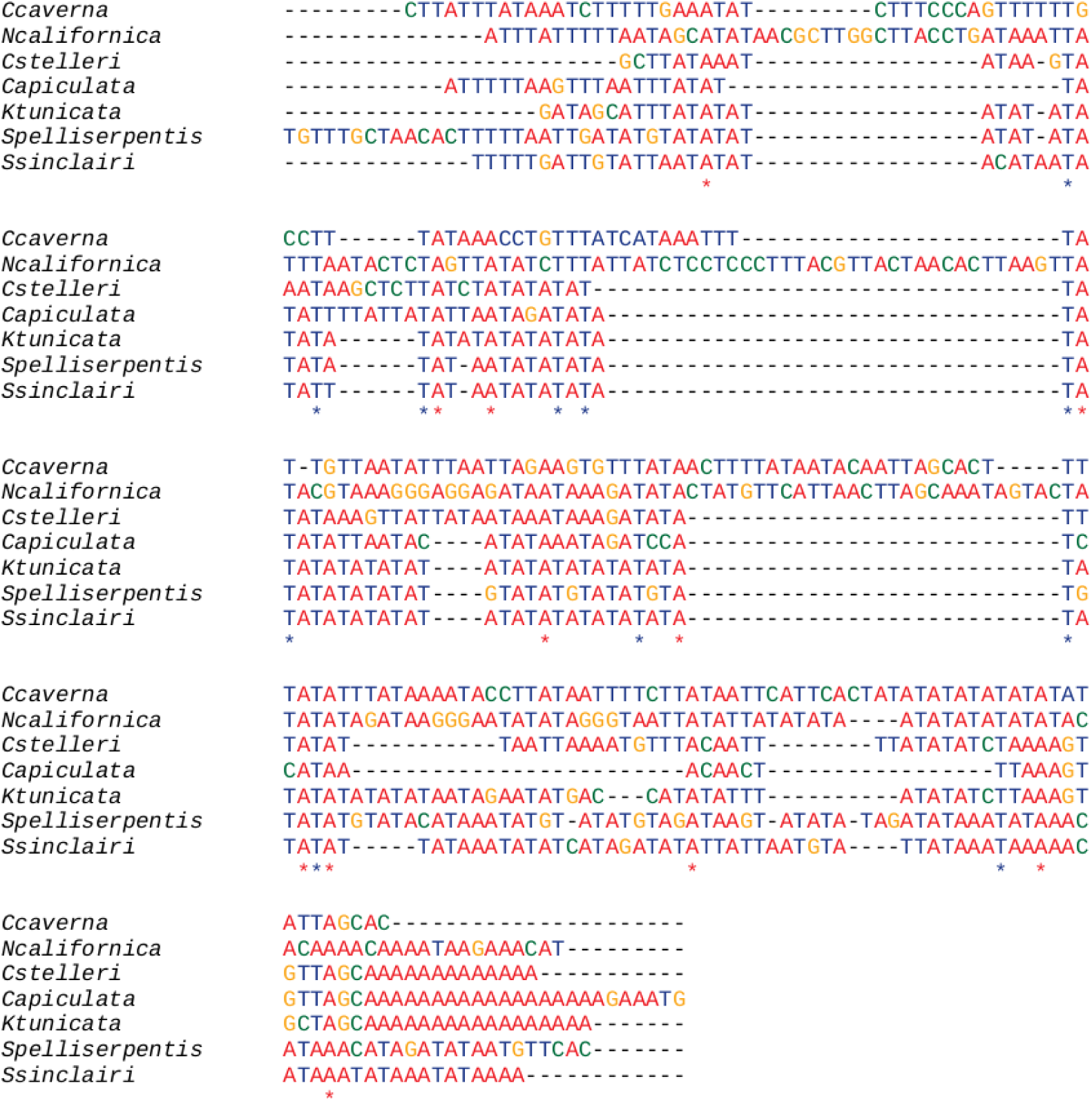
Multiple sequence alignment of the control region using MUSCLE.

**Figure 5:**
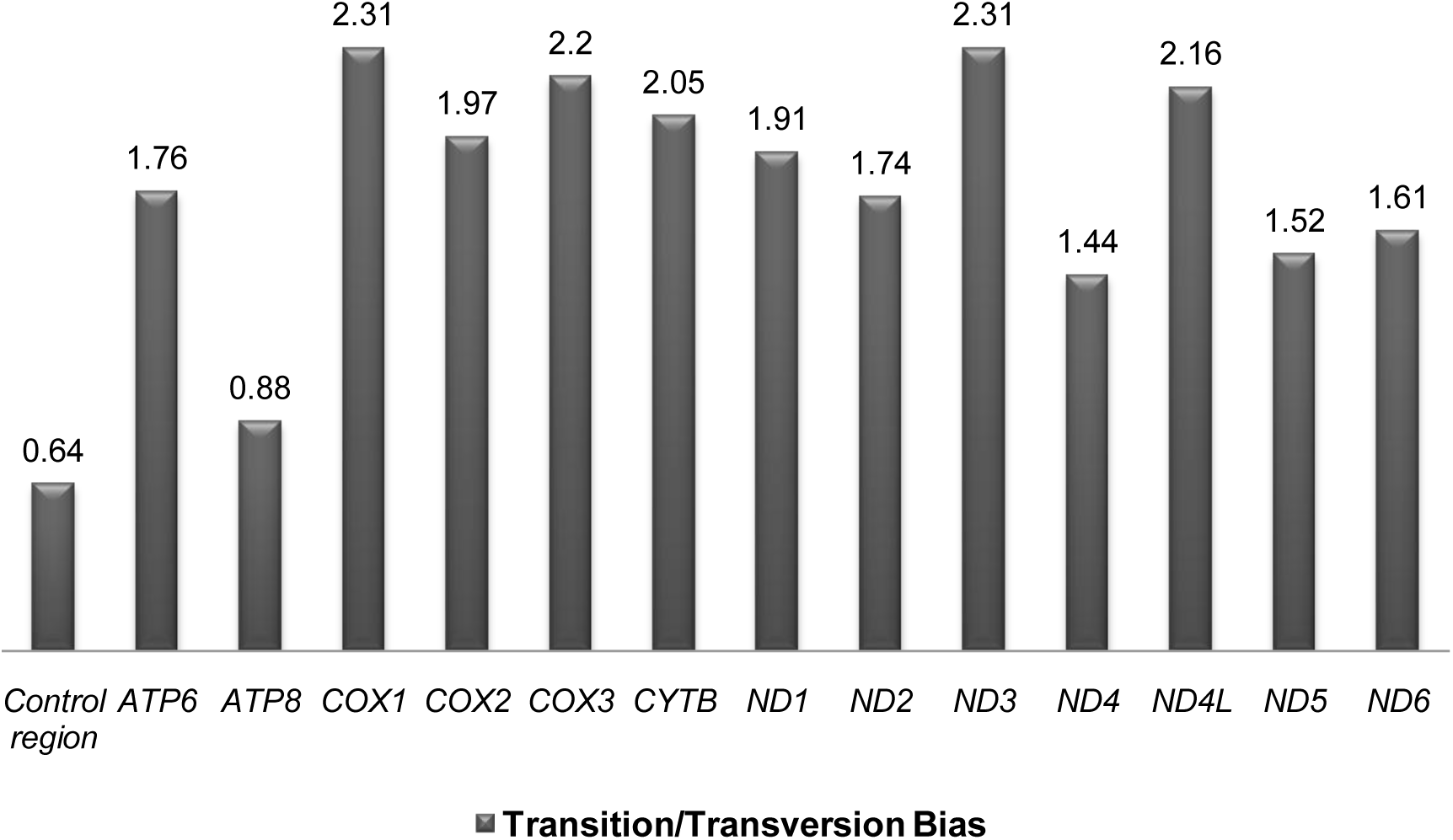
Transition *vs*. transversion ratio for the 13 PCGs and control region of the chiton mitogenomes.

### Selection studies

Purifying selection is the prevalent force in the evolution of mitogenomes. Since mitochondria serves as the major site for energy metabolism and production in the cell, weak and/or episodic positive selection may act on its protein-coding genes allowing adaptation to changes in the surrounding environment **[68, 69]**. Marine molluscs constitute ecologically important organisms in the intertidal zone and have successfully acclimatized to a dynamic environment between the tides. Considering the dynamic habitat of intertidal zone which may influence the function of mitochondrial genes, we examined signatures of positive selection in the polyplacophoran species using the CODEML algorithm implemented in EasyCodeML.

The heterogeneous selective pressure among sites in each of the 13 mitochondrial PCGs was tested, using different site-specific models: M0 (one-ratio), M1a (nearly neutral), M2a (positive selection), M3 (discrete), M7 (beta), M8 (beta and ω > 1) and M8a (beta and ω = 1). According to the results obtained, purifying selection dominated the evolution of all mitogenes except for *ATP8* **(Supp. Table 3)**. In other words, likelihood ratio test of M7 *vs*. M8 found an evidence of positive selection for only *ATP8* at codon position of 44 (LRT *p*-value = 0.000204865 and ω = 36.08464). The outcomes of comparing different site models (M0 *vs*.M3, M1a *vs*. M2a, M7 *vs*. M8, and M8a *vs*. M8) revealed the global molecular evolutionary rates of mitochondrial PCGs in the polyplacophorans under negative constraints. Furthermore, FEL analyses revealed that purifying selection was prevalent on all mitochondrial PCGs, with few sites evolving under neutrality (*p*<0.05). According to the FEL results, complex IV subunits (*COX1, COX2* and *COX3*) exhibited the highest percentage of codons under negative selection **(Fig. 6)**. *CYTB* from complex III also displayed high levels of purifying selection whereas members of complex I and V showed comparatively more relaxed selection. Additionally, we identified nine positively selected sites in six genes (*COX1, CYTB, ND2, ND3, ND4* and *ND5*) using MEME (*p*<0.05) which employs a mixed-effects maximum likelihood approach to identify individual sites subjected to episodic diversifying selection **(Supp. Table 4)**.

**Figure 6:**
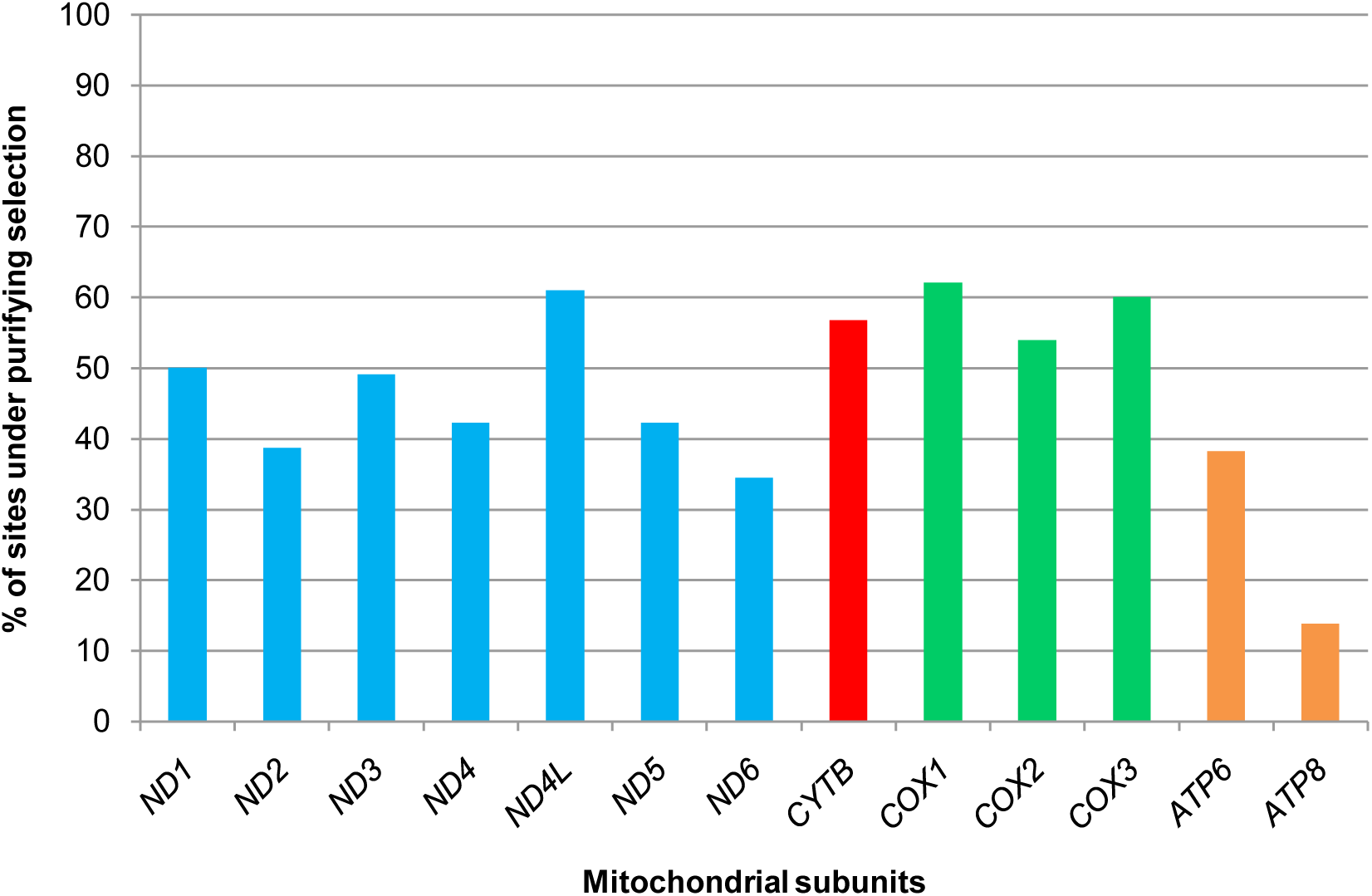
Percentage of sites under purifying selection in chiton mitochondrial subunits, detected by FEL test. The subunits are colored according to their corresponding complexes: complex I (blue), complex III (red), complex IV (green) and complex V (orange).

Since positive selection occurs at only few positions in a particular lineage for most genes, detection of natural selection becomes limited based on branch and site models separately. Therefore, we applied the more stringent branch-site model, which allows ω value to vary between branches and sites simultaneously, to measure divergent selective pressure on the specified foreground branch against the remaining background lineages. Using the Bayes empirical Bayes method, we detected 69 sites under potential positive selection in 12 PCGs with posterior probabilities ≥ 95% **(Fig. 7; Supp. Table 5)**. From the results, it is evident that complex I and V subunits possessed higher number of positively selected sites per branch compared to complex IV members.

**Figure 7:**
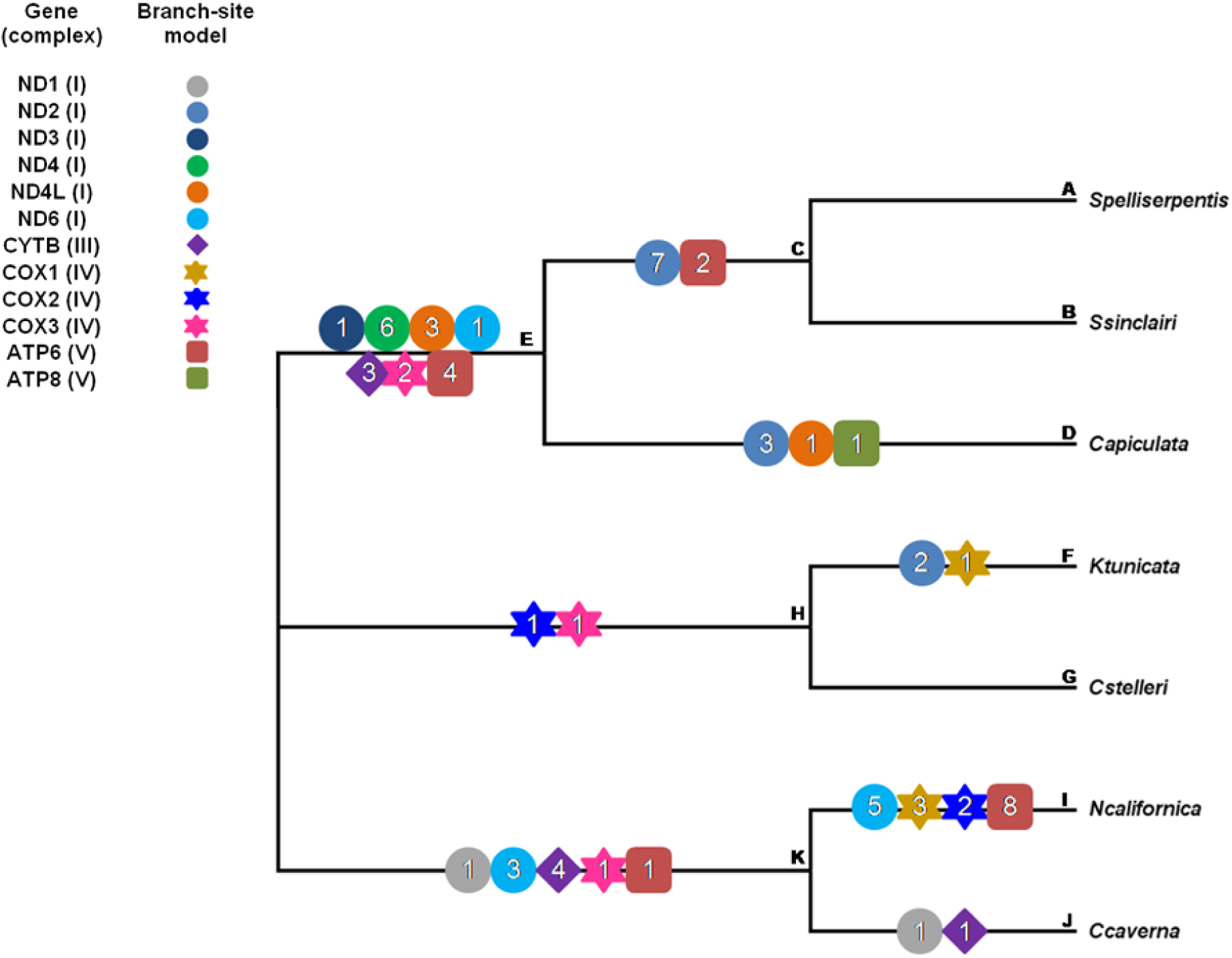
Working topology for analyzing selection pressures on each foreground branches, using the branch-site model in CODEML. The different branches of the phylogeny are labeled from A-K. Different colored shapes represent 13 mitochondrial PCGs. The number within each shape indicates the number of positively selected sites.

### Alterations of amino acid properties

The significance of ω>1 preferably indicates that positive selection would have been the driving force behind adaptive evolution of protein-coding DNA sequence. This presumption, ideally, has been found to be mostly conventional since PCGs will almost have more synonymous than nonsynonymous substitutions invariably, even if several sites have undergone positive selection pressure. Therefore, considerable changes in quantitative amino acid properties among protein residues in a phylogenetic tree were identified using the TreeSAAP software package, which compares random distributions of amino acid changes based on variable physiochemical properties with expected distributions of changes deviating from neutral conditions.

We identified several codon sites (at *p*<0.001 and category 6-8) exhibiting deviations in amino acid properties within 13 mitochondrial PCGs from seven chitons. Among these, most of the amino acid substitutions were restricted to changes in the ionization properties (*pK’*), except for *ATP8* and *ND6*. Changes in solvent accessible reduction ratio (*R*_*a*_), buriedness (*B*_*r*_), power to be at the C-terminal (*α*_*c*_) and α-helical tendencies (*P*_*α*_) were observed for *ATP6, COX1, ND1, ND2, ND4, ND5*; *ATP8, ND4, ND5*; *COX2, ND4, ND6* and *COX3, ND6*, respectively. Furthermore, *ND3* and *ND4* exhibited non-synonymous substitutions at few of their codons corresponding to changes in power to be at the middle of α-helix (*α*_*m*_) and surrounding hydrophobicity (*H*_*p*_), respectively. At the biochemical level, most of the amino acid changes in these genes affected the ionization properties while few of the substitutions influenced the hydrophobicity or hydrophilicity of amino acids. Therefore, changes in these functional properties might disrupt protein’s stability, solubility and structural organization. The alterations resulting from biochemical changes were analyzed by mapping these residues in the tertiary structures of mitochondrial protein complexes.

### Mapping of positively selected sites in mitochondrial protein structures

The putative positively selected sites, identified by CODEML and MEME, were mapped on the corresponding mitochondrial subunits, to determine their positions and significance in protein structure and function. Additionally, sites displaying changes in physicochemical properties of amino acids inferred by TREESAAP were marked **(Fig. 8a-e)**. We delineated the functional domains of individual mitochondrial subunits using TMHMM and found that the majority of sites were localized within or in the vicinity of transmembrane region. It is worth mentioning that the subunits of complex I possess more evidence of positive selection in the proton translocation channel compared to other complexes **(Fig. 9)**. There is also a clear distinction in the distribution of sites undergoing positive selection and/or radical amino acid changes between loop regions and transmembrane helices for all mitochondrial subunits.

**Figure 8:**
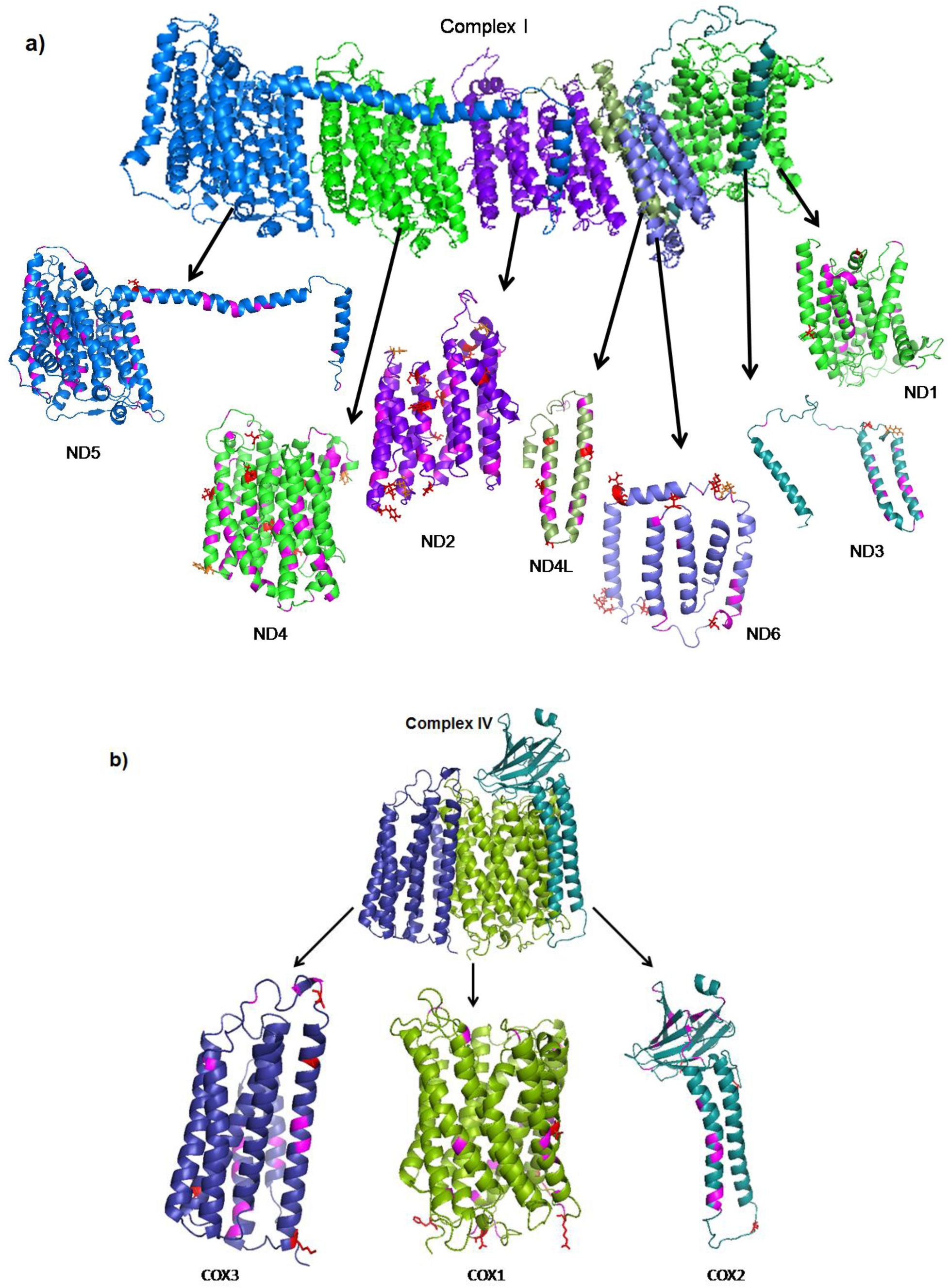
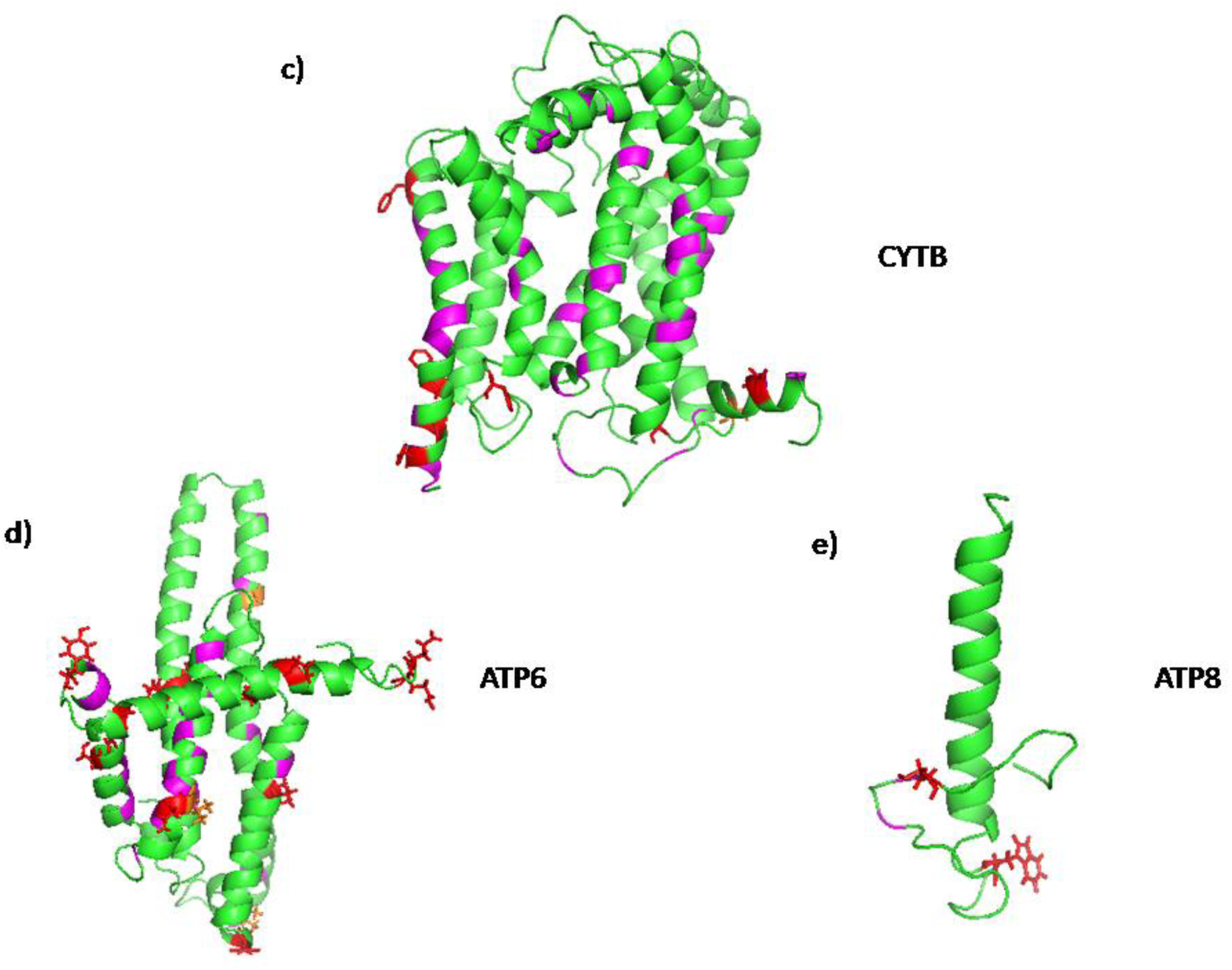
Atomic level structures of the computationally predicted mitochondrial (a) complex I, (b) complex IV, (c) CYTB subunit, (d) ATP6 subunit and (e) ATP8 subunit of *C. apiculata*. Positively selected sites inferred by CODEML and MEME are shown as red sticks whereas residues with amino acid substitutions revealed by TreeSAAP are marked in pink. Common sites identified by these methodologies are displayed as orange sticks.

**Figure 9:**
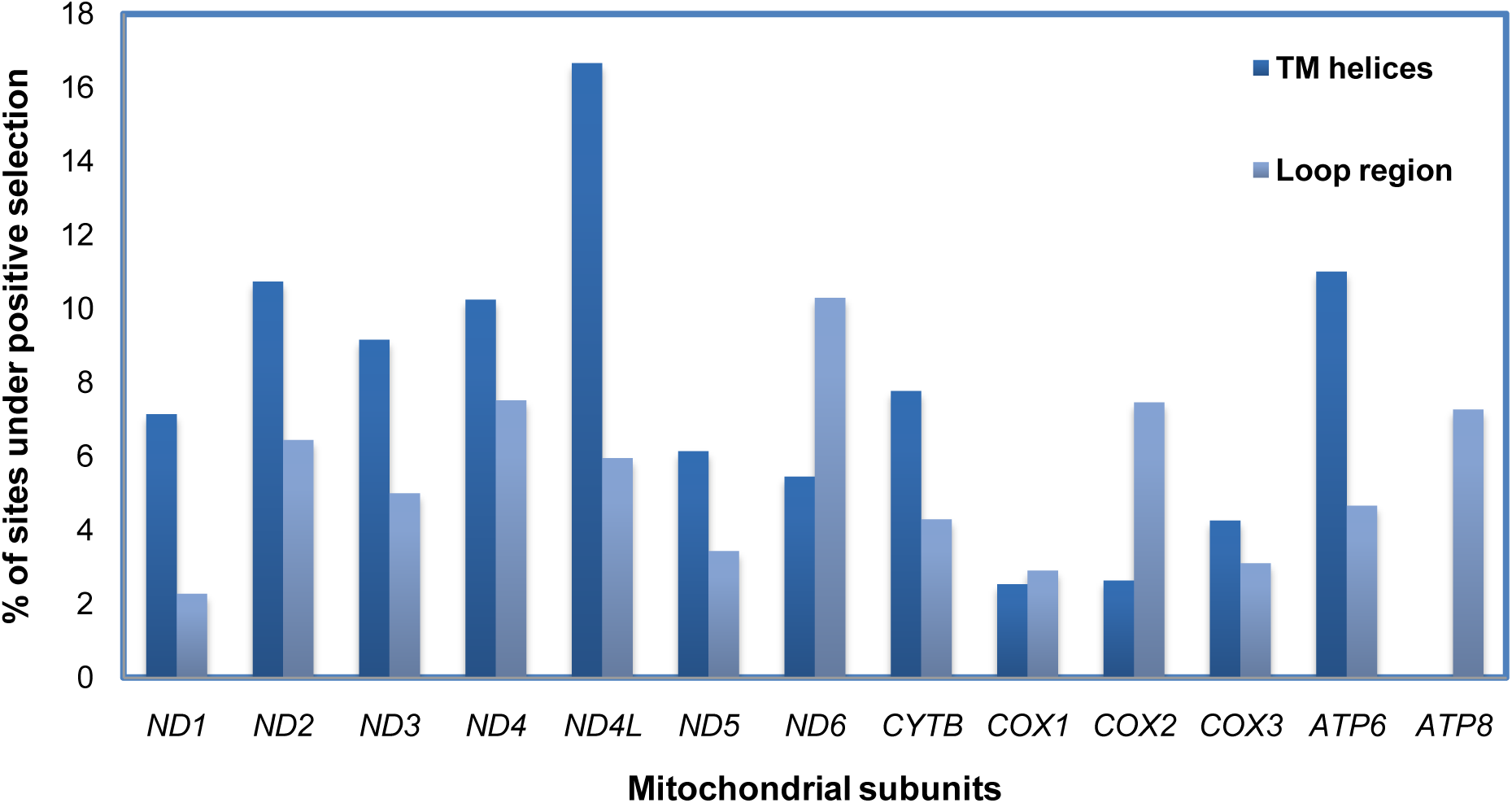
Distribution of sites, under positive selection and/or showing radical amino acid changes, between the transmembrane helices and the loop areas in individual mitochondrial subunits.

Complex I (proton-pumping NADH: ubiquinone oxidoreductase), the largest and most complicated enzyme set of the OXPHOS pathway, is responsible for NADH oxidation, ubiquinone reduction and proton pumping leading to ATP synthesis **[70]**. ND2, ND4 and ND5 are considered to be the pivotal members for maintaining proton pump due to their sequence homology with a class of Na^+^/H^+^ antiporters **[71]**. Non-synonymous substitutions in these subunits may interrupt efficacy of the proton pump. These could happen through biochemical changes that enhance/impair the proton translocation mechanism or improve/interrupt the redox linked conformational changes **[72]**. As described by de Fonseca *et al*. 2008, the high number of sites with radical amino acid changes observed in ND proteins can be attributed to their overall high mutation rate.

*CYTB*, a conserved protein constituting complex III of the electron transport chain, plays a fundamental role in energy production by catalyzing reversible electron transfer from ubiquinol to cytochrome *c* coupled to proton translocation **[73]**. We observed instances of positive selection in the functionally significant domains of *CYTB*, which may be correlated with the adaptation of chitons to the fluctuating environment of the intertidal zone. In concordance with previous literatures **[74, 75]**, we got substantial evidence of positive selection in the *COX* subunits (*COX1, COX2* and *COX3*) which is indicative of adaptive tolerance in intertidal organisms to intermittent hypoxia. Furthermore, sites under diversifying selection were identified in members of ATP synthase, the final enzyme complex of the respiratory chain that is coupled with the electrochemical gradient across the inner mitochondrial membrane to drive ATP production **[76-78]**. Therefore, the mutations in *ATPase* genes provide implications on adaptive evolution of the stress-tolerant intertidal chitons.

Intertidal molluscs display elevated sensitivity to environmental stressors like thermal stress, changing ion concentration, fluctuation in osmolarity and reduced oxygen availability. Besides, these marine invertebrates exhibit striking resilience to hydrogen sulphide (H_2_S) toxicity, a potent mitochondrial poison that disrupts mitochondrial respiratory chain by inhibiting cytochrome *c* oxidase **[79, 80]**. At this juncture, survival of these organisms relies on a modified and adaptive energy metabolism. Our results revealed potential signatures of positive selection on mitochondrial PCGs of chitons. The study provides valuable insights into the molecular mechanisms underlying the adaptive strategies of these polyplacophorans to the intertidal habitat from a mitochondrial perspective.

## Conclusion

Our study represents the first thorough investigation of evolutionary selection acting on the mitochondrial protein-coding genes of Polyplacophorans. Signatures of both positive and negative selection identified, may provide significant implications towards the adaptive evolution of mitogenomes in chitons inhabiting the intertidal zone. Additionally, mapping the positively selected sites in tertiary protein structures of OXPHOS subunits revealed their presence in functional domains, predominantly in the proton translocation channel. In conclusion, our findings provide compelling evidence of positive selection in the mitochondrial PCGs, suggestive of their role in metabolic adaptations to contrasting environments.

## Abbreviations

mtDNA: mitochondrial DNA
OXPHOS: oxidative phosphorylation
PCGs: protein-coding genes
*r*RNA: ribosomal RNA
*t*RNA: transfer RNA
ATP: adenosine triphosphate
NAD: nicotinamide adenine dinucleotide
ND: NADH dehydrogenase
CytB: cytochrome *b*
COX: cytochrome *c* oxidase complex
ATPase: ATP synthase.

