## Supplementary files for "Understanding the adaptive evolution of mitochondrial genomes in intertidal chitons"

**Supplementary Table 1:** Details of the chitons considered for this study.

| Species | Family | Accession no. | Length of mitogenome (bp) | Reference |
|---|---|---|---|---|
| *Chaetopleura apiculata* | Chaetopleuridae | KY824658.1 | 15108 | Guerra *et al.* (2018) |
| *Sypharochiton pelliserpentis* | Chitonidae | NC_024174.1 | 15048 | Veale *et al.* (2016) |
| *Sypharochiton sinclairi* | Chitonidae | NC_024173.1 | 15028 | Veale *et al.* (2016) |
| *Nuttallina californica* | Lepidochitonidae | NC_026849.1 | 15604 | Irisarri *et al.* (2014) |
| *Cyanoplax caverna* | Lepidochitonidae | NC_026848.1 | 15141 | Irisarri *et al.* (2014) |
| *Cryptochiton stelleri* | Mopallidae | NC_026850.1 | 15082 | Irisarri *et al.* (2014) |
| *Katharina tunicata* | Mopallidae | NC_001636.1 | 15532 | Boore & Brown (1994) |

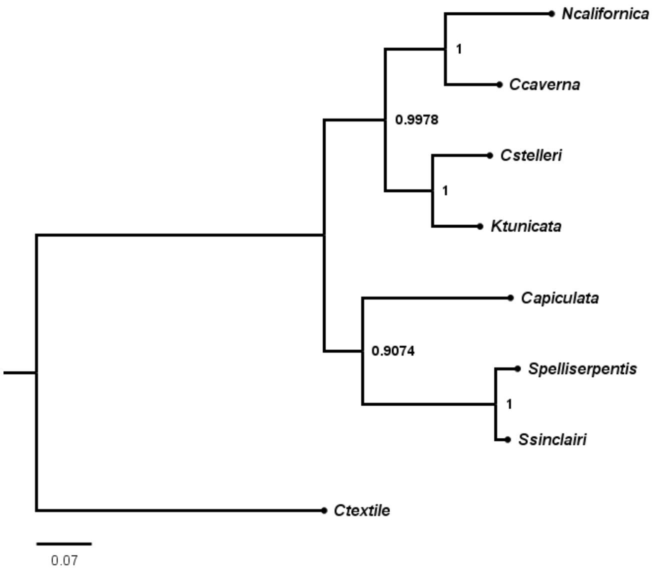

**Supplementary Figure 1(a):** Bayesian inference based topology considering the concatenated sequences of 22 *t*RNAs from seven chiton species. *C. textile* was considered as outgroup. Posterior probabilities supporting each node are shown.

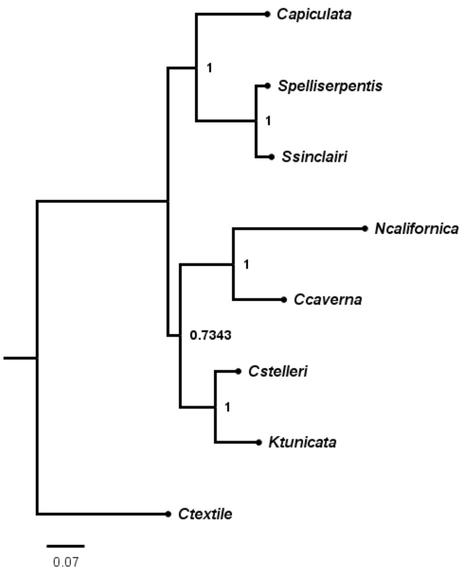

**Supplementary Figure 1(b):** Bayesian inference based topology considering the concatenated sequences of 2 *r*RNAs from seven chiton species. *C. textile* was considered as outgroup. Posterior probabilities supporting each node are shown.

**Supplementary Table 2:** Maximum likelihood estimation of substitution matrix for the PCGs and control region, performed in MEGA X.

| Gene | | A | T/U | C | G | Maximum Log likelihood values | Transition/ Transversion bias (*R*) |
|---|---|---|---|---|---|---|---|
| | **A** | - | 8.4471 | 1.4851 | 8.9869 | | |
| *ATP6* | **T/U** | 5.3211 | - | 9.7381 | 0.4467 | | |
| | **C** | 2.9596 | 30.8071 | - | 6.5331 | | |
| | **G** | 17.5074 | 1.3813 | 6.3865 | - | -4056.328 | 1.76 |
| | **A** | - | 5.7616 | 0.7890 | 6.9773 | | |
| *ATP8* | **T/U** | 5.2532 | - | 4.2702 | 1.9488 | | |
| | **C** | 2.8138 | 16.7031 | - | 16.4915 | | |
| | **G** | 19.8018 | 6.0661 | 13.1235 | - | -943.411 | 0.88 |
| | **A** | - | 9.4783 | 0.3599 | 12.7148 | | |
| *COX1* | **T/U** | 6.7430 | - | 11.6927 | 0.3194 | | |
| | **C** | 0.6785 | 30.9853 | - | 4.3864 | | |
| | **G** | 18.5846 | 0.6563 | 3.4009 | - | -6730.098 | 2.31 |
| | **A** | - | 7.7683 | 1.4731 | 13.7791 | | |
| *COX2* | **T/U** | 6.2163 | - | 9.0071 | 1.4976 | | |
| | **C** | 3.0500 | 23.3042 | - | 3.2941 | | |
| | **G** | 24.4630 | 3.3226 | 2.8246 | - | -3565.918 | 1.97 |
| | **A** | - | 9.1835 | 0.4367 | 9.7142 | | |
| *COX3* | **T/U** | 6.1940 | - | 13.9930 | 0.7229 | | |
| | **C** | 0.7229 | 34.3391 | - | 5.2427 | | |
| | **G** | 13.5416 | 1.4941 | 4.4154 | - | -3934.733 | 2.2 |
| | **A** | - | 9.2473 | 0.6861 | 12.3739 | | |
| *CYTB* | **T/U** | 5.6800 | - | 9.9234 | 0.3648 | | |
| | **C** | 1.2110 | 28.5160 | - | 5.9777 | | |
| | **G** | 19.6867 | 0.9449 | 5.3882 | - | -5808.975 | 2.05 |
| | **A** | - | 8.5160 | 0.1382 | 13.1572 | | |
| *ND1* | **T/U** | 5.4976 | - | 7.8497 | 0.4702 | | |
| | **C** | 0.3189 | 28.0544 | - | 8.8868 | | |
| | **G** | 20.1105 | 1.1132 | 5.8873 | - | -5106.734 | 1.91 |
| | **A** | - | 6.7396 | 0.3950 | 12.6499 | | |
| *ND2* | **T/U** | 4.5341 | - | 6.2770 | 0.8149 | | |
| | **C** | 1.0274 | 24.2681 | - | 11.2149 | | |
| | **G** | 22.3297 | 2.1382 | 7.6112 | - | -6274.564 | 1.74 |
| | **A** | - | 5.4323 | 1.4245 | 12.0959 | | |
| *ND3* | **T/U** | 3.7018 | - | 8.4612 | 2.0365 | | |
| | **C** | 3.7018 | 32.2656 | - | 2.0365 | | |
| | **G** | 21.9871 | 5.4323 | 1.4245 | - | -2004.887 | 2.31 |

| | | | | | | | |
|---|---|---|---|---|---|---|---|
| *ND4* | **A** | - | 8.9710 | 1.0285 | 9.1929 | | |
| | **T/U** | 6.2599 | - | 7.5851 | 0.6881 | | |
| | **C** | 2.6529 | 28.0394 | - | 7.6904 | | |
| | **G** | 19.4834 | 2.0899 | 6.3187 | - | -7699.487 | 1.44 |
| *ND4L* | **A** | - | 7.4335 | 0.0000 | 13.5261 | | |
| | **T/U** | 4.8089 | - | 8.2303 | 0.9202 | | |
| | **C** | 0.0000 | 28.2273 | - | 7.2741 | | |
| | **G** | 21.9509 | 2.3083 | 5.3205 | - | -1754.143 | 2.16 |
| *ND5* | **A** | - | 9.1139 | 1.0479 | 8.5433 | | |
| | **T/U** | 6.7343 | - | 9.0822 | 0.5663 | | |
| | **C** | 2.5484 | 29.8917 | - | 6.6793 | | |
| | **G** | 18.2773 | 1.6397 | 5.8757 | - | -9929.746 | 1.52 |
| *ND6* | **A** | - | 5.4443 | 3.4123 | 10.8099 | | |
| | **T/U** | 3.2699 | - | 5.7792 | 0.9638 | | |
| | **C** | 9.4435 | 26.6298 | - | 6.1603 | | |
| | **G** | 20.7382 | 3.0785 | 4.2703 | - | -3128.537 | 1.61 |
| *Control Region* | **A** | - | 9.2996 | 1.5578 | 4.0583 | | |
| | **T/U** | 9.2996 | - | 4.0583 | 1.5578 | | |
| | **C** | 9.2996 | 24.2270 | - | 1.5578 | | |
| | **G** | 24.2270 | 9.2996 | 1.5578 | - | -1125.298 | 0.64 |

**Supplementary Table 3:** Log-likelihood values of the site-specific models (CODEML) for *ATP8*.

| **Model** | **No. of parameters (np)** | **LnL** | **Model compared** | **LRT *p*-value** | **Positive Sites** |
|---|---|---|---|---|---|
| M3 | 17 | -837.143002 | M0 vs. M3 | 0.000000000 | - |
| M0 | 13 | -881.828094 | | | Not allowed |
| M2a | 16 | -853.115959 | M1a vs. M2a | 1.000000000 | - |
| M1a | 14 | -853.115959 | | | Not allowed |
| M8 | 16 | -838.583856 | M7 vs. M8 | 0.000204865 | 22 M 0.948, 44 C 0.967* |
| M7 | 14 | -847.077014 | | | Not allowed |
| M8a | 15 | -842.279835 | M8a vs. M8 | 0.006551613 | Not allowed |

**Supplementary Table 4:** Genes and sites under positive selection, detected by MEME in the HyPhy package.

| **Gene** | **Sites under positive selection (*p*-value <0.05)** |
|---|---|
| *COX1* | 507 |
| *CYTB* | 14 |
| *ND2* | 74, 225 |
| *ND3* | 117 |
| *ND4* | 2, 300, 389 |
| *ND5* | 482 |

**Supplementary Table 5:** Parameter estimates and likelihood values for PCGs inferred using the branch-site model. Sites under positive selection are marked with * at $p < 0.05$ & ** at $p < 0.01$

| Gene | Branch | Models (Number of parameters) | lnL | Likelihood Ratio Test (LRT) *p*-value | Positively selected sites |
|---|---|---|---|---|---|
| *ATP6* | C | Model A (16)<br>Model A Null (15) | -3520.395410<br>-3520.806490 | 0.364549389 | 94 S 0.998**<br>105 L 0.977* |
| | E | Model A (16)<br>Model A Null (15) | -3515.87977<br>-3518.721861 | 0.017118518 | 8 Q 0.987*<br>33 Q 0.968*<br>50 S 0.962*<br>124 S 0.965* |
| | I | Model A (16)<br>Model A Null (15) | -3505.435324<br>-3508.751060 | 0.010019243 | 7 E 0.962*<br>17 P 0.980*<br>39 Y 0.984*<br>70 V 0.953*<br>121 S 0.976*<br>124 S 0.955*<br>170 G 0.952*<br>207 L 0.978* |
| | K | Model A (16)<br>Model A Null (15) | -3526.295771<br>-3526.295771 | 1.000000000 | 69 N 0.952* |
| *ATP8* | D | Model A (16)<br>Model A Null (15) | -850.602983<br>-850.602983 | 1.000000000 | 30 W 0.952* |
| *COX1* | F | Model A (16)<br>Model A Null (15) | -5762.277719<br>-5764.268241 | 0.046015035 | 499 H 0.988* |
| | I | Model A (16)<br>Model A Null (15) | -5751.066253<br>-5753.626005 | 0.023658378 | 327 R 0.971*<br>404 R 0.970*<br>491 D 0.976* |
| *COX2* | H | Model A (16)<br>Model A Null (15) | -3103.062964<br>-3105.449412 | 0.028911154 | 49 K 0.998** |
| | I | Model A (16)<br>Model A Null (15) | -3102.698834<br>-3102.736834 | 0.782793109 | 6 Q 0.983*<br>205 S 0.982* |
| *COX3* | E | Model A (16)<br>Model A Null (15) | -3415.190076<br>-3419.147264 | 0.004904349 | 106 T 0.951*<br>217 I 0.997** |
| | H | Model A (16)<br>Model A Null (15) | -3418.700076<br>-3419.807070 | 0.136764722 | 71 K 0.978* |
| | K | Model A (16)<br>Model A Null (15) | -3415.056594<br>-3415.278312 | 0.505468690 | 97 A 0.968* |
| *ND1* | J | Model A (16)<br>Model A Null (15) | -4411.035287<br>-4411.559978 | 0.305649461 | 287 S 0.985* |
| | K | Model A (16)<br>Model A Null (15) | -4415.651735<br>-4415.651735 | 1.000000000 | 161 S 0.986* |
| *ND2* | C | Model A (16)<br>Model A Null (15) | -5563.703249<br>-5563.782031 | 0.691409257 | 36 N 0.981*<br>70 Y 0.974*<br>78 L 0.975*<br>108 S 0.954*<br>114 C 0.953*<br>135 S 0.974*<br>316 W 0.952* |
| | D | Model A (16)<br>Model A Null (15) | -5566.494831<br>-5566.494831 | 1.000000000 | 97 S 0.980*<br>167 S 0.979*<br>236 S 0.978* |

| | | | | | |
|---|---|---|---|---|---|
| | F | Model A (16)<br>Model A Null (15) | -5566.472186<br>-5567.956701 | 0.084872732 | 121 W 0.954*<br>215 Q 0.973* |
| *ND3* | E | Model A (16)<br>Model A Null (15) | -1805.264409<br>-1808.912759 | 0.006908138 | 114 Y 0.980* |
| *ND4* | E | Model A (16)<br>Model A Null (15) | -6950.584020<br>-6954.066838 | 0.008308989 | 46 S 0.960*<br>60 S 0.983*<br>267 K 0.969*<br>294 G 0.984*<br>322 S 0.989*<br>373 F 0.994** |
| *ND4L* | D | Model A (16)<br>Model A Null (15) | -1361.212224<br>-1361.264972 | 0.745331326 | 17 C 0.984* |
| | E | Model A (16)<br>Model A Null (15) | -1358.749310<br>-1359.768974 | 0.153277350 | 27 S 0.998**<br>38 G 0.957*<br>58 S 0.988* |
| *ND6* | E | Model A (16)<br>Model A Null (15) | -2936.774563<br>-2936.911784 | 0.600367486 | 78 S 0.967* |
| | I | Model A (16)<br>Model A Null (15) | -2935.117597<br>-2935.117597 | 1.000000000 | 44 E 0.951*<br>107 K 0.993**<br>110 M 0.982*<br>151 K 0.973*<br>154 Y 0.980* |
| | K | Model A (16)<br>Model A Null (15) | -2933.082458<br>-2934.482546 | 0.094253961 | 128 S 0.954*<br>129 L 0.994**<br>162 P 0.976* |
| *CYTB* | E | Model A (16)<br>Model A Null (15) | -5126.708470<br>-5130.743683 | 0.004499354 | 18 S 0.995**<br>361 F 0.974*<br>365 N 0.976* |
| | J | Model A (16)<br>Model A Null (15) | -5138.463278<br>-5139.330524 | 0.187838243 | 367 Y 0.970* |
| | K | Model A (16)<br>Model A Null (15) | -5122.220963<br>-5127.377406 | 0.001321045 | 8 I 0.955*<br>76 T 0.965*<br>301 H 0.999**<br>342 F 0.976* |
